# Differences in the genomic potential of soil bacterial and viral communities between urban greenspaces and natural arid soils

**DOI:** 10.1101/2025.03.13.643106

**Authors:** María Touceda-Suárez, Alise Jany Ponsero, Albert Barberán

## Abstract

Urban green spaces provide essential ecosystem services are ever more important in arid cities. However, the design and management of these greenspaces often require physicochemical transformations whose effect in the balance of the arid urban ecosystems is normally not accounted for. In this project, we leverage metagenomic data from soil microbial communities of urban greenspaces and neighboring natural areas in a city from the arid Southwestern USA (Tucson, Arizona) to understand the differences in microbial (bacterial and viral) community structure and function in urban greenspaces compared to natural arid soils. We found bacterial and viral communities to be distinct between urban greenspace and natural arid soils, viral communities showing a starker differentiation. Urban greenspace bacteria were less versatile and showed higher genetic potential for the simple carbohydrate consumption and nitrogen reduction. Moreover, bacteria in urban greenspaces exhibit higher genetic potential for resistance to heavy metals, and certain clinical antibiotics. Our results suggest that the conversion of arid natural land to urban greenspaces determines the soil microbiome structure and functioning, and potentially its ability to adapt to the changing environment.

**IMPORTANCE:** Urban green spaces are critical for the sustainability of arid cities. Nevertheless, they require deep soil physicochemical transformations. Soil bacterial and viral communities are responsible for soil functioning and provision of some ecosystem services, but they are also highly influenced by changes in the soil environment. The significance of our research is in illustrating the structural and functional changes that microbial and viral communities undergo in urban soils of arid cities, and their potential impacts on urban greenspace soil processes.

## INTRODUCTION

Urban green spaces provide essential ecosystem services, such as cooling, water filtration, and soil stabilization, that are ever more important in arid cities (González-Méndez & Chávez-García 2020). Urban development in arid areas has been rising with rural-urban migration, and the expansion of arid lands globally (Gober & Burns 2002). Yet often, green space infrastructure in arid cities is designed according to global models of urban development which are not suited for arid conditions (Larson & Perrings 2013). For example, intensive irrigation and fertilization are needed to maintain green areas in arid cities since urban green spaces are usually made up of non-arid adapted vegetation (Pearse et al. 2016; Ignatieva et al. 2020). Thus, additionally to the disturbances posed globally by urbanization –such as air, water, and soil pollution, increased temperature, and loss of permeable surface– (Pickett et al. 2011) urban greenspaces in arid cities, and particularly their soils, often undergo complete physicochemical transformations whose effect in the balance of the urban ecosystems is not yet understood.

Soil microbial and viral communities are responsible for soil functioning and provision of some ecosystem services associated with urban greenspaces, such as soil stabilization, water regulation, carbon storage, nutrient cycling, and antibiotic resistance regulation (Ooi et al. 2022; Fan et al. 2023). At the same time, soil microorganisms are influenced by the changes in environmental conditions posed by urban development (Nugent & Allison 2022; Cleavenger et al. 2023). In fact, urbanization has been shown to, through management and pollution, homogenize soil properties (Ryan et al. 2022) and microbial community composition (Barberán et al. 2015; Epp Schmidt et al. 2017; Liu et al. 2022), and increase microbial nutrient cycling potential, and the proportion of genes associated with greenhouse emissions and resistance to xenobiotics (Delgado-Baquerizo et al. 2021). Previous studies reported increased levels of soil inorganic nitrogen, N_2_O emissions, moisture, and organic matter in urbanized arid soils (Davies & Hall 2010; Hall et al. 2008), which have been associated with changes in microbial community composition and functioning (Hall et al. 2009), and more specifically, with an increased abundance of microbial groups involved in aromatic compound degradation and ammonia oxidation (Chen, Martinez, et al. 2021). Although these studies provide first insights into the impacts of urbanization on arid soil microbial communities, further high-resolution research is needed to unveil the specific microbial functions that are modified by urbanization and their potential impact in ecosystem services’ provision. This is especially important in arid environments, wherein we know very little about the effects of urbanization on the structure and genetic potential of the soil microbiome.

Whole genome shotgun analyses (i.e., metagenomics) allow us to study the function of the soil microbiome and identify organisms without the need for prior isolation in pure culture (Fierer 2017). Importantly, assembled metagenomic contigs can be mined to identify genomic sequences from DNA bacteriophages –virus that infect bacteria– (Williamson et al. 2017). Bacteriophages, or “phages”, shape microbial community structure, and modify nutrient cycling by releasing microbial necromass (Trubl et al. 2018; Albright et al. 2022; Wieczynski et al. 2023). Additionally, phages can incorporate host genes into their genomes, serving as vectors for horizontal gene transfer upon further infections of different hosts. Although most of these genes, termed auxiliary metabolic genes (AMGs), are used by the virus to regulate the host’s metabolism and increase their own survival, some have functions, such as transformation of carbon, or degradation of organic pollutants, that could provide the bacterial host with an advantage to changes in the environment (Sun et al. 2023). On the other hand, previous studies have found soil phages to be a more responsive fraction of the soil microbial community to rapid changes in the soil environment (Santos-Medellín et al. 2023). Thus, we can expect the soil transformations driven by urbanization to influence the phage community, and those changes to influence the bacterial community through viral-host dynamics.

Here, we leverage the metagenomic information from soil microbial communities of urban greenspaces and neighboring natural areas in a city from the arid Southwestern USA (Tucson, Arizona) aiming to: (i) compare the structure and life history strategies of bacterial and bacteriophage communities between urban and natural soils; (ii) assess changes that the urban environment can cause on microbial function, such as modifications of the genetic potential for carbon and nitrogen cycling, and resistance to heavy metals and antibiotics. We hypothesize that: 1) the structure of both bacterial and phage communities would differ between urban greenspaces and natural arid soils, but phages –given their higher responsiveness to changes in the environment (Santos-Medellín et al. 2023)– would show more drastic structural differences; 2) virus in urban greenspace soils would show a preference for virulent lifestyle facilitated by higher water content caused by soil management (Liao et al. 2022); 3) microbial communities of urban greenspace soils would specialize in certain sources of carbon and nitrogen originated from urban-specific inputs of these elements (i.e., fertilization and atmospheric pollution); and 4) urban greenspace soils would show evidence of higher levels of anthropogenic contaminants, such as heavy metals and antibiotics, through a higher abundance of resistance genes.

## MATERIALS AND METHODS

### Study sites, soil sampling, and physicochemical analyses

Soil samples were collected in Tucson, AZ, USA and surrounding natural areas in August of 2019 (Supplementary Figure 1). Six samples were collected in two urban parks, Reid Park (RP 32°12’28.08’’ N, 110°55’24.96’’ W, elevation 765 m), and Himmel Park (HP; 32°14’0.42’’ N, 110°56’5.48’’ W, elevation 752 m). Twelve samples were collected in natural areas representative of the ecosystem types surrounding the city: Sonoran desert (Sabino Canyon; SC; 32°18’37’’ N, 110°49’16’’ W, elevation 830 m), ponderosa pine forest (Rose Canyon; RC; 32°23’15’’ N, 110°42’40’’ W, elevation 2,119 m), and arid shrubland (Santa Rita Experimental Range: range 11B, 31°46’25’’ N, 110°52’42’’ W, elevation 1,156 m; range 8, 31°46’30’’ N, 110°51’41’’ W, elevation 1,211 m; Exclosure45, Exc. 45; 31°48’60’’ N, 110°51’42’’ W, elevation 1,553 m; and UAB, 31°49’41’’ N, 110°50’46’’ W, elevation 1,158 m). At each site, 9 m transects were established. Surface soil samples were collected at three equidistant locations along the transect at 3 m, 6 m and 9 m, and 0–20 cm depth. Soil samples were immediately transported in sealed and sterilized plastic bags on ice to the laboratory. Soil samples were sieved through a 2-mm mesh and homogenized. Visible living plant material, like pieces of leaves and roots, was removed. A subsample was placed into a sterilized tube and stored at −80 °C until DNA extraction. The remaining soil was used for physicochemical analyses. Soil physicochemical properties: pH, Electrical Conductivity (EC), Calcium (Ca), Magnesium (Mg), Sodium (Na), Potassium (K), Nitrate (NO_3-_), Phosphate (PO_4_^3-^), exchangeable sodium percentage (ESP) and cation exchange capacity (CEC) (meq/100g); were measured from water saturated soil paste extracts produced from 200.0 ± 0.5 g of air-dry pulverized soil by Motzz Laboratory (Phoenix, AZ, USA) using the standard methods for soils of the Western region.

### Metagenomic sequencing and assembly

Total soil genomic DNA was extracted using a DNeasy PowerLyzer PowerSoil Kit (Qiagen) following manufacturers’ instructions. Genomic DNA was fragmented to 500 bp length and ligated to Illumina adapters using the QIAseq FX DNA Library Kit (Qiagen, Hilden, Germany). Quality and quantity were determined with Agilent 4150 TapeStation DNA bioanalyzer. Samples were shotgun-sequenced on a 2 × 150 bp Illumina NextSeq550 platform at the Microbiome Core of the University of Arizona. Quality of raw reads was evaluated using FastQC v.0.11.9 (Andrews 2010) and used to estimate of the parameters for adapter removal adapters with bbduck (Bushnell 2022)and quality filtering with Trimmomatic v. 0.38 (Bolger et al. 2014) (reads shorter than 50 bp and low-quality bases were removed). A total of 69,333,926 to 356,676,854 reads per sample remained after quality trimming. Reads from each sample were *de novo* assembled using MEGAHIT v. 1.1.4 (Li et al. 2015), with the k-mer length increasing from 21 to 141 in steps of 20. The contig N50 ranged from 715 to 964 bp among samples. Protein coding sequences were inferred from contigs using Prodigal v. 2.6 (Hyatt et al. 2010), and dereplicated by clustering by ≥95% similarity and ≥80% overlap using MMseqs2 v.13.45111 (Steinegger & Söding 2017). This resulted in a catalog containing 32,930,898 non-redundant genes. Paired-end reads of each sample were mapped to the gene catalog using BWA v. 0.7.16 (Li & Durbin 2009) and coverM (Aroney et al. 2024) to obtain their abundance per sample.

### Bacterial taxonomic and functional annotations

Bacterial taxonomy was annotated from clean reads using kraken2 v.1.1.1 (Wood & Salzberg 2014) against the 10/23 RefSeq standard database, followed by bracken v.3 (Lu et al. 2017) to produce a count table of bacterial species. Non-bacterial species were filtered out of this count table, as well as bacterial species with a prevalence of less than 10% of the samples, and an abundance of less than 0.00001.

We calculated life-history traits from urban and natural soil microorganisms. First, average genome size (Barberán et al. 2014), and 16S rRNA copy number were calculated following the methods described in Pereira-Flores et al. 2019. Average genome size was calculated as the average number of base pairs divided by number of genomes per sample –estimated as the mean coverage of the 35 single-copy genes–, while the average 16S rRNA copy number was estimated as the coverage of 16S rRNA genes divided by the number of genomes. Next, we calculated the GC content and the variance of GC content from community quality-filtered reads (Barberán et al. 2012). Codon usage bias –the preference of a genome for a specific set of synonymous codons, which correlates with growth rate– was calculated from ribosomal genes as the inverse mean the effective number of codons (ENC′) of all ribosomal genes (Vieira-Silva & Rocha 2010). Finally, we used annotations obtained by comparing the gene catalog against the Kyoto Encyclopedia of Genes and Genomes (KEGG) database using hmmsearch v.3.3.2 (Johnson et al. 2010), to estimate the proportion of unannotated genes and the sugar-acid preference. The proportion of unannotated genes was calculated by comparing the relative abundance of unannotated genes to that one of genes with annotations. Sugar-acid preference –which informs of the preferred source of energy of the cell, sugars or amino acids– was calculated as the weighted (SAP) and unweighted ratio between the relative abundance of genes in sugar decomposition pathways and that of genes in amino acid decomposition pathways (Gralka et al. 2023).

To annotate genes involved in the cycling of carbon and nitrogen, heavy metal resistance and antibiotic resistance in our bacterial populations, clean reads (quality-filtered and trimmed) were aligned against the CAZy v.3.2.1 (Cantarel et al. 2009), NCyc (Tu et al. 2019), BacMet v.2.0 (Pal et al. 2014) and CARD v.3.2.8 (McArthur et al. 2013) protein databases respectively using the blastx algorithm (McGinnis & Madden 2004) through diamond (Buchfink et al. 2015) with an e-value cutoff of 10e-9. Gene count tables were manually normalized to Reads Per Kilobase per Million mapped reads (RPKM). To obtain a more precise picture of differences in community C acquisition strategy, the RPKM normalized abundances of CAZy gene annotations were aggregated based on their targeted substrates (cellulose, chitin, glucan, lignin, peptidoglycan, starch/glycogen, xylan, other animal polysaccharides, other plant polysaccharides, oligosaccharides) using a curated database (Piton et al. 2023). In order to obtain the same level of precision in the differences in N cycling processes, RPKM normalized abundances of NCyc annotated genes were aggregated on the base of process (i.e., nitrification, denitrification, nitrogen fixation, etc) using a curated database (Tu et al. 2019). Genes specific to the denitrification pathway were identified from the literature for a detailed visualization of their abundances (Yang et al. 2017). BacMet annotations were aggregated based on the heavy metal they provide the cell with resistance to using the BacMet internal classification. CARD annotations were aggregated by class of the drug they provide resistance to, and resistance mechanism using the CARD ARO ontology.

### Bacteriophage inference and annotation

Viral contigs were recovered from metagenomic assembled contigs using Deepvirfinder2 (Ren et al. 2020) and Virsorter2 v.2.2.3 (Guo et al. 2021) with a minimum length cutoff of 1500 bp. Viral contigs inferred by both tools were merged and their quality and completion was assessed using CheckV v1.0.1 tool, and database v.1.4 (Nayfach et al. 2021). Viral operational taxonomic units (vOTUs) were estimated by clustering viral contigs using a 95% identity and 70% completeness cutoff (Li et al. 2022) with MMseqs2 (Steinegger & Söding 2017). Clean reads were mapped to vOTUs to obtain a vOTU count table, that was subsequently normalized to reads per base per million (RPKM). Viral reads accounted for an average of 6.4% of total metagenomic reads. A summary of the number and length of inferred viral contigs can be found in zenodo (See data availability). Taxonomy of viral species was obtained using geNomad v.1.8. (Camargo et al. 2023). In addition, life history classification of viruses in temperate vs. virulent viruses was performed using PhaTYP v.3 (Shang et al. 2023). Finally, potential host taxonomy was predicted from viral sequences using the iPHoP v1.3.3 suite (Shang et al. 2021) and virus-host ratios (VHR) were calculated as the ratio per sample between the relative abundances of the viruses and the bacterial species identified as their hosts and present in the sample. We annotated both auxiliary metabolic genes (AMGs), and antibiotic resistance genes (ARGs) in viral sequences. AMGs where inferred from viral contigs - preprocessed with virsorter2’s “prep-for-dramv” module-using Dram-v (Shaffer et al. 2020). While to annotate ARGs present in viral contigs, species representative sequences inferred from viral contigs were aligned against the CARD (McArthur et al. 2013) protein database, using the blastx algorithm (McGinnis & Madden 2004) through DIAMOND v.2.0.9 (Buchfink et al. 2015) with an e-value cutoff of 10^-9^ and a best match filter.

### Metagenome assembled genomes (MAGs) recovery and annotation

Contigs obtained from singular assembly were binned and bins were refined into MAGs using Maxbin, MetaBAT, and Concoct through the metaWRAP suite v.1.3.2 (Uritskiy et al. 2018). A total of 1,133 MAGs were recovered, we retained 66 medium to high-quality MAGs (i.e. > 70% completion, <10% contamination) for downstream analyses (Bowers et al. 2017). MAG taxonomy was annotated using the GTDB-tk v.3 (Chaumeil et al. 2020) database and tool. MAG abundance per sample was calculated by mapping quality cleaned reads from back to the MAG collection using BWA v. 0.7.16 (Li & Durbin 2009) and CoverM v.0.6.1 (Aroney et al. 2024). to create a MAG count table. Metabolic gene annotation was performed using DRAM v.3 (Shaffer et al. 2020). In order to locate viral sequences in bacterial MAGs, viral species sequences were aligned against a database constructed from medium-high quality MAGs using BLAST with a min e-value cut off of 10e-9. Multiple matches of the same viral and bacterial sequence were filtered to conserve only the best match. Antibiotic resistance genes and heavy metal resistance genes were located in MAGs by aligning of the MAG sequences against the CARD and BAcmet databases using blast (McGinnis & Madden 2004) with a min e-value cut off of 10e-9. Multiple matches of the same viral and bacterial sequence were filtered to conserve only the best match.

### Statistical analyses

Statistical analyses were implemented in R v4.2.2 (R Core Development Team 2015). To evaluate the effect of urbanization on soil microbial richness, number of different genes, and relative abundance of genes, we used a mixed-effects model with site as a random effect using *lme4* (Bates et al. 2015). We implemented a PERMANOVA on each dataset to test the effect of urbanization controlling for the effect of site on microbial community composition and genetic profile composition using the *vegan* package v.2.6-4 (Oksanen et al. 2020). Differential abundance of annotated genes and abovementioned classifications were calculated using the Maaslin2 package v.1.12.0 including a fixed effect of urbanization with a site as a random (Mallick et al. 2021). To evaluate the association between virus and their inferred hosts present in our samples, Pearson’s correlation coefficients were calculated between the relative abundance per sample of each virus and said bacterial host.

## RESULTS

### Urban greenspace soils present distinct microbial communities, bacteria with lower genome sizes, and more virulent viruses

We found 3,572 bacterial species on average per sample, and 2,306 vOTUs on average per sample (Figure 1). The number of different species (i.e., richness) did not differ significantly between soils from urban greenspaces and natural soils for either bacteria (X^2^ = 0.67, p-value = 0.412; Figure 1A) or phages (X^2^ = 0.72, p-value = 0.397; Figure 1C), but viral abundance was slightly higher in natural soils (X^2^ = 25.02, p-value < 0.001; Supplementary Figure 1). Additionally, when comparing among natural sites representing different habitats, we saw that phage richness was lowest in ponderosa pine forests with respect to all other soils –urban and natural grassland, and desert shrubland– (Supplementary Figure 1).

**Figure 1.**
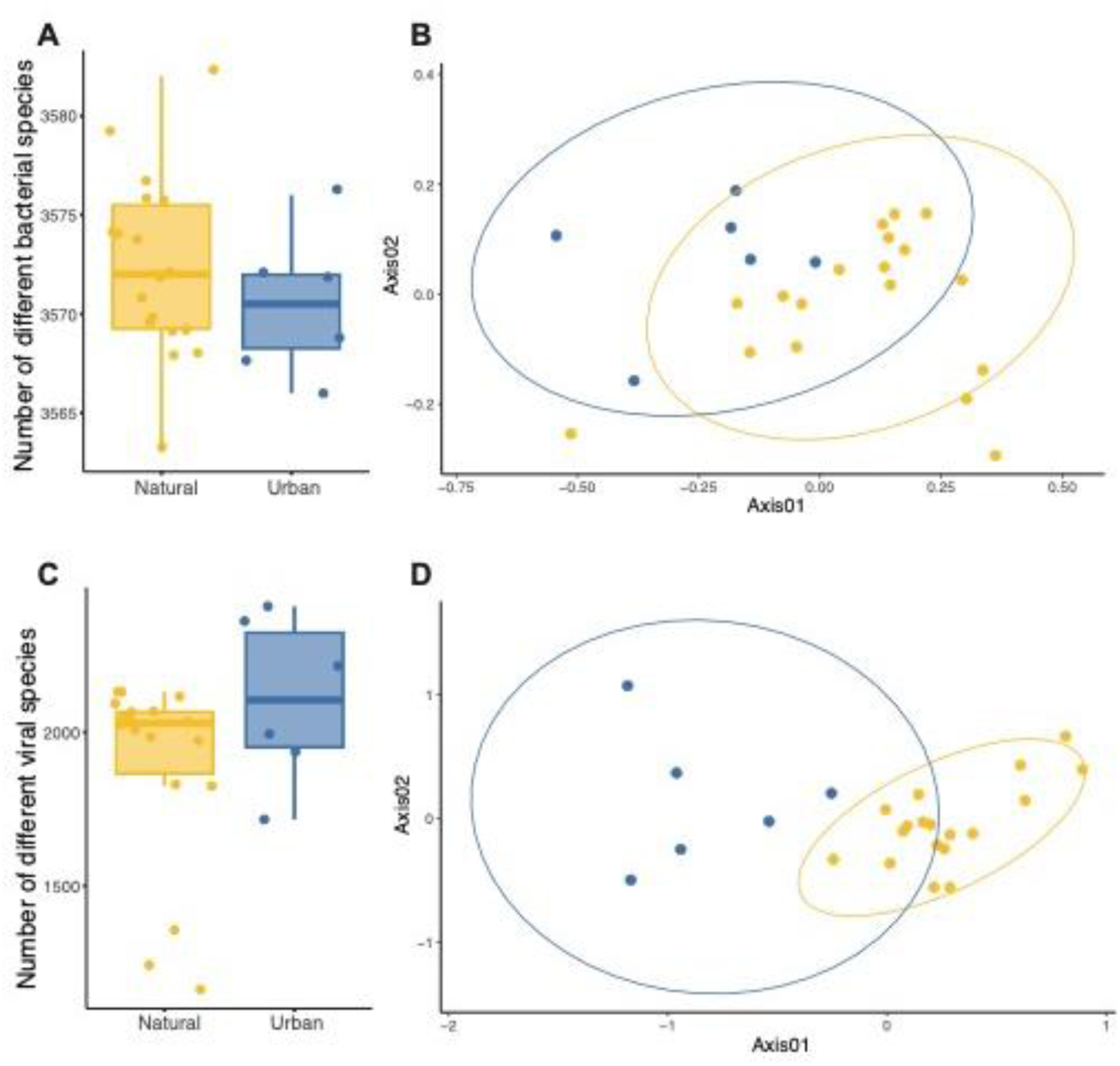
Community structure of soil bacteria and phages in urban greenspaces and natural soils. A and. **C**, Number of different bacterial and viral species in urban greenspace (blue) and natural (yellow) soils. **B and D,** Bacterial and viral community similarity respectively.

Both bacterial and phage community compositions were different in urban compared to natural soils (bacteria: R^2^ = 0.20, p-value = 0.001; phages: R^2^ = 0.24, p-value = 0.001; Figure 1B and D). Nevertheless, we found great overlap between all sites, both urban and natural, for both bacteria and viruses, and urban sites had a higher number of distinct vOTUs, but not bacteria (Supplementary Figure 2 and Supplementary table 1). Accordingly, soil physicochemical composition was also different between urban and natural soils, especially for micronutrients such as Ca, NO_3-_, and Na, exchangeable sodium potential (ESP), and cation exchange capacity (CEC; Supplementary Figure 3). Some of the bacterial families whose abundance changed more dramatically from natural to urban greenspace soils were Geodermatophilaceae and Nocardiodaceae (increased abundance in urban soils), and Mycobacteriaceae and Bradyrhizobiaceae (reduced abundance in urban soils; Supplementary Figure 4). Additionally, microbial community composition also differed between natural habitats around Tucson, the most different bacterial and phage communities being those found in ponderosa pine forests (Supplementary Figure 1).

In addition to changes in community composition, we found differences in metagenomics-resolved life history traits between urban greenspace and natural soils. Bacteria in urban soils presented lower average genome sizes than in natural soils (X^2^ = 33.73, p-value < 0.001; Figure 2A), but higher average 16S rRNA copy number (X^2^ = 7.19, p-value = 0.007; Figure 2B), GC content (X^2^ = 2.15, p-value = 0.145; Figure 2C), and proportion of unannotated genes (Supplementary Figure 4). The GC content variance, however, was lower in urban soils (X^2^ = 3.688, p-value = 0.055; Supplementary Figure 4), while no differences were found in codon usage bias (X^2^ = 0.33, p-value = 0.565; Supplementary Figure 4). In terms of the characterization of viral communities, only 13% of viral genomes were annotated taxonomically–most of them classified as undetermined Caudoviricetes– which is not sufficient to make any meaningful comparisons (Supplementary Figure 5). However, we were able to predict the lifestyle of 39% of the inferred viral contigs and compare their abundance in urban vs. natural soils. We found that while abundance of temperate phages was higher than that of virulent phages in both soils, (urban: F = 0.9, p-value = 0.364; natural: F = 16.77, p-value < 0.001; Figure 2D), the ratio of virulent/temperate virus abundance was slightly higher in urban soils (urban = 0.67, natural = 0.54).

**Figure 2.**
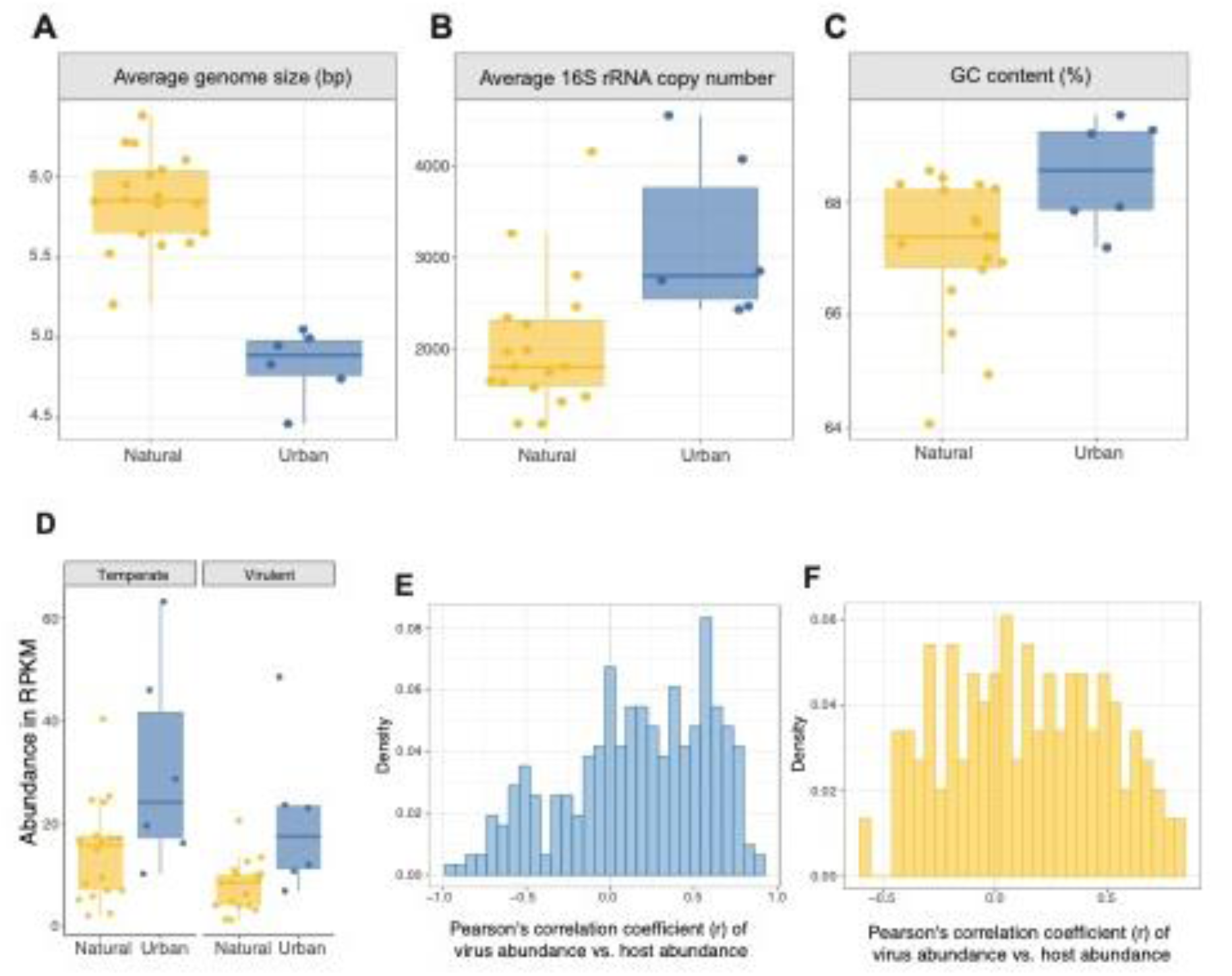
Functional traits of bacterial and viral communities in urban greesnpaces and natural soils. **A-C**, Metagenomics-informed bacterial life history traits. **D,** Relative abundance –in reads per kilobase per million mapped reads (RPKM)– of inferred viral lifestyles. **E-F,** Density distribution of Pearson correlation coefficient values between virus abundance (RPKM) and predicted host abundance (RPKM).

### All soils have higher proportion of lysogenic viruses, but carbon utilization AMGs are more abundant in urban greenspaces

We predicted the potential hosts of 23% of the inferred viral contigs. Although most high abundance hosts were shared in both urban and natural soils (i.e., Geodermatophilaceae, Streptomycetaceae, Rubrobacteraceae), urban soils showed a uniquely high abundance of certain families, such as Nitrososphaeraceae and Coleofasciculaceae, and reduced abundance of Acetobacteraceae compared to natural soils (Supplementary Figure 5). We calculated the Pearson’s correlation coefficient (*r*) between the abundance of each phage and that of its host (Ma et al. 2024). In urban soils, the frequency distributions of *r* were highly skewed to the left, values between 0.5 and 1 being the most frequent; while in natural soils the distribution of *r* values was multimodal, showing peaks around *r* = 0, and *r* = [0.25-1] (Figure 2E and F). Correlation values closer to +1 could suggest a higher proportion of lysogenic viruses (in a 1 to 1 abundance relationship with their host do to mutualism), while values closer to -1 would signify a higher proportion of lysogeny (1 to - 1 abundance relationship with their host due to predation). Furthermore, we calculated the Pearson’s correlation coefficient between the virus-host abundance ratio (VHR) and the host abundance, which follows the opposite logic to the correlation between abundances. Most correlation values for pairs in both urban and natural soils were negative, while pairs in urban soils showed slightly higher frequencies of positive values (Supplementary Figure 6). Additionally, when plotting the bacterial genus abundance per sample against the number of putative links with viruses present in said sample we found, in both urban and natural soils, that our observations clustered around three areas of the plot: low host abundance, low viral presence (most of the observations); low host abundance, medium to high viral presence; and medium to high host abundance low viral presence; but did not find many cases of bacterial hosts having both high abundance and high viral presence (Supplementary Figure 6).

A total of 18 putative auxiliary metabolic genes (AMGs) were identified from the bacteriophage contigs, 16 of them from bacteriophages inferred from urban soils. Of these 16 genes, five of them were involved in organic nitrogen utilization, and one of them was involved in carbon utilization (Supplementary table 2). Both AMG groups showed higher abundance in urban soils (Supplementary Figure 7). No genes of antibiotic resistance were found in viral genomes.

### Urban greenspace soil bacteria harbor less diversity of carbon and nitrogen cycling genes, and are specialized on simpler carbohydrates decomposition and denitrification

Although the abundance of C cycling genes was similar in urban and natural soils (X^2^ = 0.97, p-value = 0.325; Figure 3A) and across different natural habitats (Supplementary Figure 8), natural soils harbored a higher C cycling gene richness –measured as the number of different C cycling genes– (X^2^ = 10.777, p-value = 0.001; Figure 3B) with ponderosa pine forest having the highest values of richness and shrubland the lowest among natural soils (Supplementary Figure 8). Furthermore, by comparing the abundance of genes in different pathways, we observed that urban soil microorganisms had a higher abundance of genes involved in the metabolism of simple starch and glycogen substrates, while natural soils had higher abundance of genes involved in the metabolism of more complex compounds (i.e., dextran, glucan, chitin, cellulose; Figure 3E).

**Figure 3.**
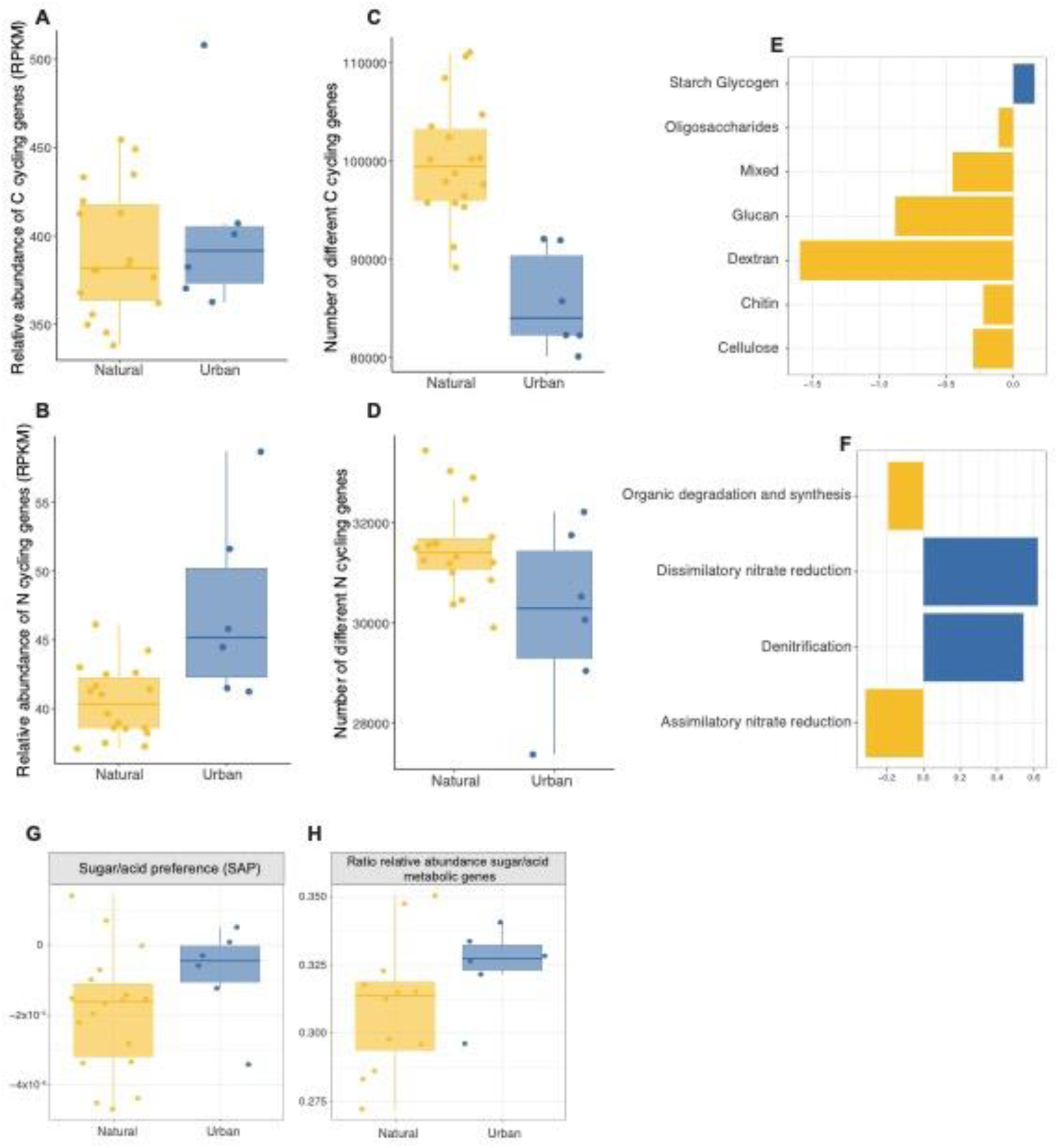
Carbon and nitrogen cycling potential of soil microbial communities under urbanization. **A-B** Relative abundance of genes involved in C and N metabolism. **C-D,** Number of different genes involved in C and N metabolism. **E,** Differential abundance of C cycling genes in pathways of decomposition of different carbohydrates**. F,** Differential abundance of N cycling genes involved in different processes within the N cycle. **G-H,** Weighted (SAP) and unweighted ratio between the relative abundance of genes involved in sugar metabolism and genes involved in acid metabolism.

Urban soils had higher relative abundance of nitrogen cycling genes (X^2^ = 7.27, p-value = 0.007; Figure 3B) and, similarly to the C cycling genes, lower richness (X^2^ = 3.81, p-value = 0.051; Figure 4D), with forest soils showing the lowest values of richness among natural habitats (Supplementary Figure 8). Specifically, urban soils showed a higher abundance of genes involved in denitrification (conversion of oxidize nitrogen forms to nitrogen gas) and dissimilatory nitrate reduction (reduction of nitrate in the absence of oxygen; Figure 3F). All genes in the denitrification pathway except nirS were more abundant in urban soils (i.e., narG, nirK, norB, nosZ; Supplementary Figure 8). Additionally, none of the genes involved in ammonia oxidation had higher abundances in urban soils (Supplementary Figure 8). Finally, urban soils had higher values of both weighted sugar-acid ratio, or SAP, (X^2^ = 0.81, p-value = 0.367; Figure 3G), and unweighted ratio of relative abundance of sugar decomposition genes vs. acid decomposition genes (X^2^ = 0.88, p-value = 0.349; Figure 3H).

**Figure 4.**
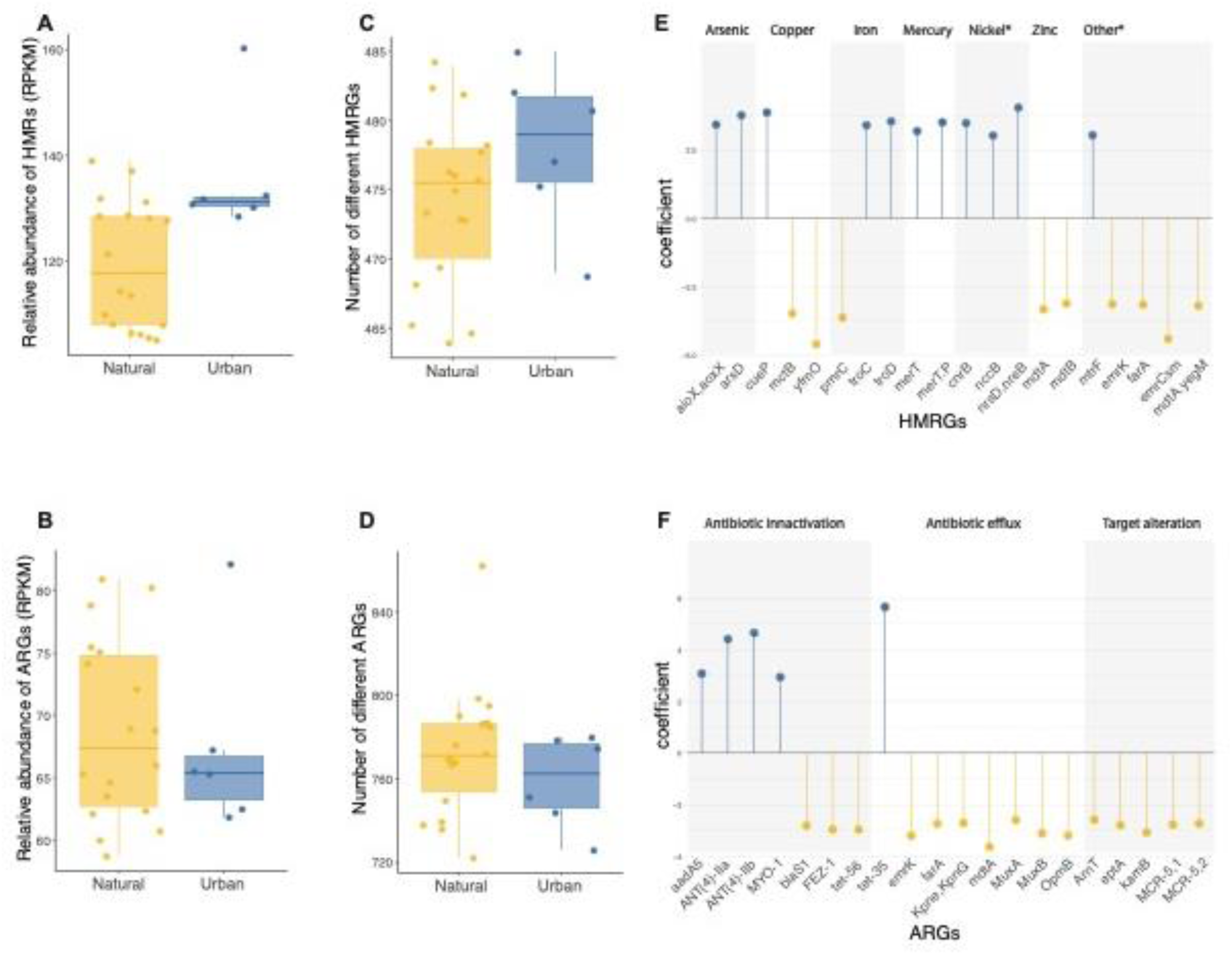
Potential for heavy metal and antibiotic resistance in urban greenspace and natural soils. **A-B** Relative abundance of HMRGs and ARGs. **C-D,** Number of different HMRGs and ARGs. **E-F,** Differential abundance of HMRGs and ARGs.

### Urban greenspace soil bacteria have a higher load of heavy metal but not antibiotic resistance genes

Both the relative abundance of heavy metal resistance genes (HMRGs) and their richness were higher in urban soils compared to natural soils (Relative abundance: X^2^ = 8.0688 p-value = 0.030; Richness: X^2^ = 11.611 p-value = 0.003; Figure 4A and C), with no differences between natural habitats (Supplementary Figure 9). Differential abundance calculations on resistance genes grouped by heavy metal or compound resulted on 51 groups at a 0.2 q-value cutoff level (Supplementary Figure 10), eight of them being more abundant in urban soils (i.e., silver, antimony, gold, mercury, manganese, copper, arsenic, and cobalt). This translated into 222 genes differentially abundant at a 0.2 q-value threshold. When selecting the top 20 most differentially abundant genes –genes with highest absolute differential abundance coefficient value– urban soils presented higher abundance of genes of resistance to arsenic, iron, mercury, and nickel, while resistance genes to copper and zinc were more abundant in natural soils (Figure 4E).

Contrary to the HMRGs, neither the abundance nor the richness of antibiotic resistance genes (ARGs) were significantly different in urban vs. natural soils (Relative abundance: X^2^ = 0.12, p-value = 0.733; Richness: X^2^ = 1.73, p-value = 0.188; Figure 4B and D). Only one resistance mechanism (i.e., antibiotic target replacement) was differentially abundant between natural and urban soils using a q-value > 0.2 threshold, being higher in urban soils (Supplementary Figure 10). However, 15 drug classes were differentially abundant over that same 0.2 q-value threshold, seven of them having higher abundance in urban soils (i.e., pleuromutilin, panem, penam, streptogramin A, cephamycin, oxazolidinone, and macrolide; Supplementary Figure 10). These groups translated in 168 differentially abundant ARGs. Out of the top 20, five were more abundant in urban soils: aminoglycoside nucleotidyltransferase genes aadA5, ANT(4’)lla and ANT(4’)llb, and myosin gene MYO1, all involved in antibiotic inactivation, and tetracycline efflux pump gene tet35, which is involved in antibiotic efflux (Figure 4F).

### Metagenome assembled genomes (MAGs) do not reflect the changes in the community

We reconstructed 66 high quality MAGs. Out of those 66 MAGs, 44 were reconstructed from natural samples, while 22 were reconstructed from urban green space samples. Most of the MAGs belonged to the phyla Actinomyceota, Acidobacteriota, Chloroflexota, and Cyanobacteriota (Supplementary Figure 11). We did not find any specific metabolic pathway involved in carbon or nitrogen cycling to be enriched in MAGs highly present in urban samples (Supplementary Figure 12). Three out of the 66 MAGs carried a HMRG, two carried an ARG, and four carried both. Despite the trends in abundance and richness of HMRGs in the community, MAGs carrying HMRGs were less abundant in urban samples (Supplementary Figure 12). Out of the 66 MAGs, 44 of them had one or more sequence that aligned with the inferred viral sequences –a prophage sequence– (Supplementary Figure 12). Finally, the proportion of base pairs belonging to prophage sequences in the MAGs recovered was negatively correlated with the MAG’s GC content in urban soils (*r* = -0.61, p-value= 0.002; Supplementary Figure 6), and less strongly in natural soils (*r* = -0.30, p-value= 0.048; Supplementary Figure 6).

## DISCUSSION

The soil modifications required to maintain greenspaces in arid climates, together with disturbances associated by the urban environment, may produce changes in the function of soil microbial communities and consequentially on ecosystems dynamics (Grimm et al. 2008). In this study, we leveraged soil metagenomes from urban greenspaces and neighboring natural ecosystems in a city from the arid Southwest US (Tucson, Arizona) to study these changes. Briefly, we started by analyzing the community structure of soil bacteria and bacteriophages and found large differences in community composition and ecologically informed bacterial functional traits –such as genome size, average rRNA copy number, and GC content– between urban greenspaces and natural soils, but we found no changes in richness for either microbial group. We then studied the virus-host associations and found most viruses to be temperate in both urban and natural soils, and their abundances to be coupled to the abundance of their hosts in most cases. Additionally, an analysis of the association between host abundance and the number of virus that could infect said host revealed three interaction types: low host abundance - low putative viral links; low host abundance –medium/high putative viral links; and medium/high host abundance– low putative viral links. Finally, we analyzed the abundances of different groups of bacterial genes involved in the processing of nutrients (carbon and nitrogen) and pollutants (heavy metals and antibiotics) enriched in or associated with urban environments. Urban greenspaces showed a specialization in simple forms of carbon, and an enhanced potential for cycling of nitrogen –specifically denitrification –, and resistance to heavy metals. Lastly, we found that, while the potential for antibiotic resistance was not higher in urban greenspace soils, we did find genes of resistance to clinically used antibiotics in these soils.

Differences in both bacterial and viral community composition between urban greenspace and natural soils were in agreement with our hypothesis that urban soil transformation leads to changes in soil microbial structure, and consistent with previous findings (Chen, Martinez, et al. 2021). However, given that the difference in bacterial community composition have been also seen in different cities around the globe (Delgado-Baquerizo et al. 2021), it seems that these changes are a consequence of the differences between urban greenspaces (i.e. management) and natural soils irrespectively of climate. Importantly, viral communities revealed less species presence overlap between urban and natural soils than bacterial communities, confirming previous evidence of high heterogeneity in viral species distribution (Santos-Medellín et al. 2023; Barnett & Shade 2024). On the other hand, we found no differences in either bacterial or viral richness between urban and natural soils. As a matter of fact, the effects of urbanization on microbial species richness have been contested in the literature (Ramirez et al. 2014; Abrego et al. 2020; Whitehead et al. 2022), which suggests that there is no uniform effect of urbanization, and that factors such as microbial group (Grierson et al. 2023), urbanization intensity (Chen, Martinez, et al. 2021), city characteristics (Aronson et al. 2014), urban site characteristics (Reese et al. 2015), and spatial scale (Concepción et al. 2015) may modify the response soil microbial communities have to the complex urban environment. Thus, our results confirm that in our Southwestern US city, urban soils harbor a bacterial and viral diversity comparable to that one of natural arid soils (Chen, Martinez, et al. 2021).

To further illustrate the functional consequences of the differences in community composition between urban greenspaces and natural arid soils, we explored community-level metagenomics-informed bacterial life history traits. The use of life history traits has been gaining attention in microbial ecology, partly because they allow to sort through the overwhelming amount of data on taxonomical groups and gene abundances associated with metagenomics studies, and distill a classification of the microbial communities funded on their interaction with the environment (Fierer et al. 2014). For example, a recent study used several genomic traits of Sonoran Desert bare and vegetated soil microbial communities to characterize them in the copiotroph-oligotroph framework (Chen, Neilson, et al. 2021). In our study, we found bacterial communities of urban greenspace soils to have lower average genome size and higher 16S rRNA copy number, suggesting that these communities are dominated by bacteria that are less metabolically versatile, and thus less able to deal with environmental variability (Barberán et al. 2014; Westoby et al. 2021), but are in turn capable of rapidly responding to resource availability and trigger growth (Roller et al. 2016). These traits match what would be expected of soil microorganisms specialized in a homogeneous and stable environment with periodic input of nutrients and water, such as an urban park. At the same time, the reduced genomic versatility acquired through this specialization could pose a disadvantage for these microorganisms in the face of changing climatic conditions (Fitzpatrick & Keller 2015). Interestingly, average community GC content in our microbial communities was higher in urban greenspace soil communities. High GC content is associated with thermal resistance since DNA molecules with high GC content tend to be more stable (Yakovchuk et al. 2006), and thus could be a result of higher soil temperatures caused by the Urban Heat Island effect, which is poorly regulated in grass dominated urban parks (Kraemer & Kabisch 2022). Alternatively, the higher GC content could be associated with the higher abundance of Actinobacteria in these soils (Fierer et al. 2007). Together, these results reflect the adaptations we could expect from a microbial community subjected to a warmer, moister, and more nutrient rich soil environment. Unfortunately, they could also reflect a reduced adaptability to changing conditions.

On the same line, we know that viral life strategies and virus-host interactions are influenced by differences in the soil environment. For instance, increased water in soil pore space enhances the propagation of lytic viruses and, since infection requires physical contact, their ability to infect bacterial hosts (Liao et al. 2022). This could explain why we found higher levels of predicted virulent viruses and AMGs for viral reproductive success in urban soils compared to natural soils (Luo et al. 2022). However, most of the viruses we inferred from both urban and natural soils were predicted to have temperate lifestyles, as is generally the case in soils (Williamson et al. 2007). These results, together with the higher occurrence of positive correlations between the relative abundances of viruses and their hosts, the lower correlation values between soil virus-host ratio and microbial density, and the lack of differences in host targeting potential, suggests that most of the soil viral community replicates with their host (lysogeny) in both natural and urban greenspace soils –although increased moisture and temperature in urban greenspaces could cause peaks in lytic activity –, and points to a potential dominance of a Piggyback-The-Winner strategy (Knowles et al. 2016; Ma et al. 2024). It is important to note, however, that current methods used in this work offer limited annotations –we were able to annotate the lifestyle of 39%, and host of 23% of the viral community –, and so our conclusions should be revised upon improvement of the annotation software. Additionally, our viral analysis are based on metagenomics inferred viral genomes, and thus do not maximize the viral genome recovery nor do they illustrate the active viral community (Kosmopoulos et al. 2023).

The changes in bacterial community structure and life strategies were accompanied by changes in genetic potential (Fierer et al. 2012). Changes in microbial biogeochemical cycling potential of C and N between natural and urban soils are of special interest given the continuous expansion of urbanized land and its contribution to greenhouse gas emissions (Koerner & Klopatek 2002; Hall et al. 2009). Numerous studies have tackled this issue, showing that, contrary to elements that are dependent on the soil parent material (i.e., P and K), C and N are deeply influenced by anthropogenic activities (Trammell et al. 2020). For example, the C cycle can be influenced in urban soils through deposition and accumulation of inorganic carbon and black carbon from industrial and vehicle emissions, and through the increase of organic carbon from introduced vegetation and management dependent (Vasenev & Kuzyakov 2018). In our study, the effects of such increased organic matter are evidenced by the reduced diversity of C cycling genes in urban soils compared to natural soils, which could be due to a reduced variability in C sources. Furthermore, despite the general abundance of C cycling genes did not increase in urbanized soils compared to natural soils, urban soils had a preference for sugar metabolism, and higher abundance of genes involved in the degradation of starch, a plant produced polysaccharide that is usually decomposed in early stages, while natural soils presented genes, at both the community and the genome level, for the decomposition of a more diversified range of C sources, most of them being late decomposition stage polysaccharides, such as cellulose, and microbially produced polysaccharides (i.e., dextran and chitin) (Martin 1971). Additionally, the fact that this specialization has not been seen in other cities (Delgado-Baquerizo et al. 2021) supports the hypothesis that vegetation changes and management are key factors altering the C cycling potential of microbial communities in arid cities specifically (Golubiewski 2006; Trammell et al. 2020).

Simultaneously, the N cycle in urban soils is altered by the introduction of inorganic N through fertilization and atmospheric deposition (Hall et al. 2008). This is evidenced in our study by the higher abundance of N cycling genes found in urban soil microbial communities, and the decreased diversity of those genes, again suggesting a decrease in the variability of N sources. The introduction of inorganic N fertilizers has been associated with increase in the levels of nitrous oxide (N_2_O) emissions in both agricultural and urban managed soils (Ravishankara et al. 2009). N_2_O is greenhouse gas with a global warming potential 300 times that of CO_2_ whose concentrations, together with those of its reduced form of nitric oxide (NO), in the atmosphere have grown exponentially primarily due to human activity (IPCC 2019). Production of N_2_O in soils happens primarily through denitrification of nitrate (NO_3_^-^) to N_2_, which we found to be the pathway with more highly abundant genes in urban soils, together with dissimilatory nitrate reduction, another pathway of production of N_2_O (Asamoto et al. 2021). However, enrichment of denitrification pathways has been seen in urban greenspace soils in other climates (Delgado-Baquerizo et al. 2021), suggesting that the emission of this gas is not climate specific. Furthermore, our study is limited to the genetic potential of microbial communities, and our results are therefore not a good indicator of the measure of metabolization of N forms into N_2_O. Studies looking at the expression of the genes involved in the production, and consumption of N_2_O should be coupled with measuring the emission of these gases from urban soils to answer these questions.

Urban soil microorganisms are exposed to pollutants originated from human activities, which makes them a critical laboratory to study the effects this exposure could have on natural systems in the long run. Since our health is closely tied to the environmental microorganisms that we are in contact with, one very relevant question to explore in soil metagenomes is how exposure to anthropogenic pollution modifies the genetic make-up of these microbes (Li et al. 2018). Two prevalent forms of this pollution in urban environments, and specifically soils, are heavy metals and antibiotics. Both heavy metals and antibiotics can hamper the survival of soil microbes, and thus microbes will respond to the presence of these substances by encoding molecular barriers –or resistance mechanisms– in their genomes. Supporting this notion, we found the abundance and diversity of heavy metal resistance genes to be higher in urban soils compared to natural soils. Gene specific analysis revealed a higher abundance of genes providing resistance to some common urban pollutants such as copper (Cu), mercury (Hg) and arsenic (As) (Wen Liu et al. 2023), while genes of resistance to other metals such as lead (Pb) and zinq (Zn) were highly abundant in natural soils, which could be originated through the influence of contamination expanding over the city limits, or through presence of these metals in the soil’s parent material (Yu-Rong Liu et al. 2023). Surprisingly, microbes from natural soils present resistance genes to a wider variety of pollutants, including growth enhancers (i.e., hydrazine), fertilizer related pollutants (i.e., halogens), and anti-parasites (i.e., salicylanilide).

In contrast with a previous study targeting urban greenspace soils of 23 cities globally distributed (Delgado-Baquerizo et al. 2021), we did not find the abundance or richness of ARGs to be higher in urban greenspaces compared to natural soils. We hypothesize that this discrepancy could be due to the ARGs genes included in the study. In our study we included all ARGs present in the CARD database, while the other studies select clinically relevant ones. Based on this, our results suggest that the adoption of antibiotic resistance as a response to competition –evidenced by larger genome sizes– in natural soils is comparable to that one responding to human pollution with antibiotics in urban soils. Another possible explanation would be for urban pollution to reach city adjacent natural sites (Yu-Rong Liu et al. 2023). Most probably, our results are originated from a combination of both of the previous potential mechanisms; since when we interrogated further on the class of drugs microorganisms present resistance to, we find a combination of drugs of present clinical use (i.e. tetracyclin, fluoroquinolones, elfamycin, nitrofuran), and compounds that are still not developed for clinical use (i.e. free fatty acids) (Casillas-Vargas et al. 2021).

In summary, this study analyzed the differences in microbial structure and function in urban greenspaces and natural soil of an arid Southwestern US city. We found that urban greenspaces in this city harbor less versatile bacterial communities with efficient growth strategies based on simple carbohydrate sources, and elevated potential for resistance to heavy metals and certain clinical antibiotics. Furthermore, bacterial communities from urban greenspaces showed a higher potential for denitrification, which leads to the production of N_2_O, a potent greenhouse gas. Although our analysis was limited to the genetic potential of soil microorganisms and should be confirmed through measurements of soil gas emissions; these results suggest that the management required to maintain urban greenspaces in arid cities can contribute to global greenhouse emissions, as well as to the generation of soil microbial communities with a reduced resilience to changing conditions. The design of greenspaces in arid cities could focus on low management alternatives, such as native vegetation, to balance their ecosystem services and disservices. (Guillen-Cruz et al. 2021). All in all, our findings enhance our understanding of the impacts that the management of urban greenspaces has on the soil microbiome and provide evidence of the role of soil microorganisms in the shifts in ecosystem dynamics produced by the soil transformations introduced in urban greenspaces of arid cities.

## Data availability

Scripts for the analysis and visualization of data can be found at https://github.com/merytouceda/urban_greenspaces_aridcities. Final data products from processing can be found at 10.5281/zenodo.13152735. Raw metagenomics sequences are available at http://www.ncbi.nlm.nih.gov/bioproject/1143147, reference number PRJNA1143147

