## Supplemental figures for "Differences in the genomic potential of soil bacterial and viral communities between urban greenspaces and natural arid soils"

**Supplementary Information**

**Supplementary Figures**

**Supplementary figure 1. Microbial community structure based on vegetation type. A. and C.** Boxplots representing the bacterial and viral richness (respectively) in urban and natural soils of different vegetations (bacteria: X^2^ = 0.33, p-value = 0.850; virus: X^2^ = 25.14, p-value < 0.001). **B. and D.** NMDS ordination plots where points represent bacterial and viral (respectively) community composition of each soil sample, colored by vegetation type (bacteria: R^2^ = 0.20, p-value = 0.019; virus: R^2^ = 0.22, p-value < 0.001). Ellipses represent urban and natural groups from Figure 1. **E.** Boxplots of the number of predicted viral contigs with virulent or temperate lifestyles in urban and natural soils and colored based on vegetation. Only differences between vegetations of natural sites were significant (F = 7.11, p=value = 0.003).

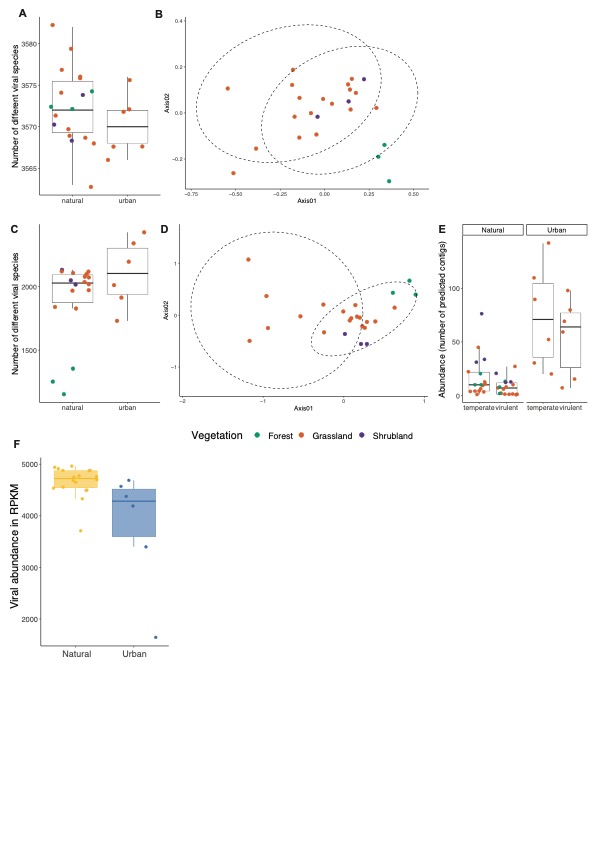

**Supplementary Figure 2.** **Bacterial and viral species overlap between soils**. A. Bacterial and B. viral species upset plots. Columns represent the number of species present in the soils marked with a solid sphere in the lower panel, or the species overlap between those soils. The size of the yellow and blue bars on the left bottom corner represent the total number of species present in the sample. Replicates have been combined by site.

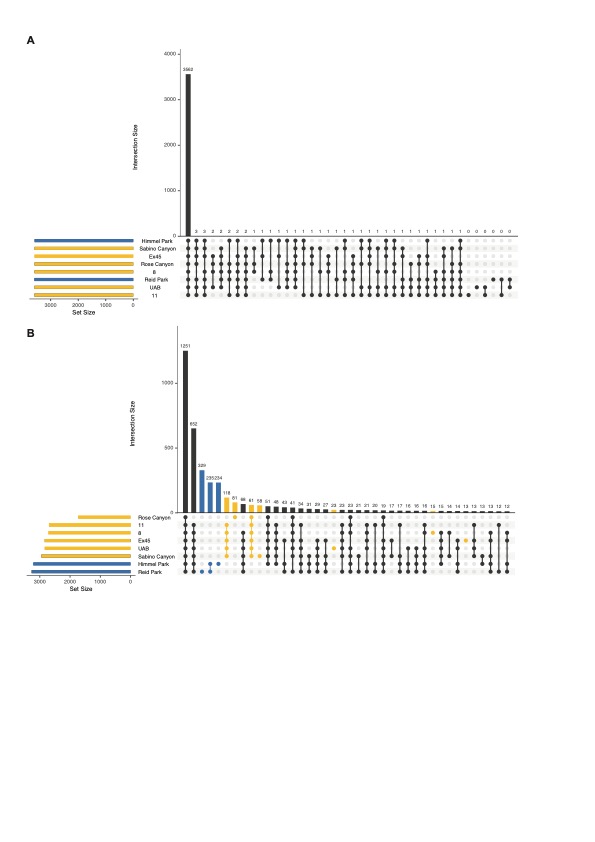

**Supplementary figure 3. Geographical distribution and physicochemical properties of soil samples. A.** Location of all 8 sampling sites around Tucson, AZ, USA. Points are yellow for natural samples, and blue for urban samples. Colors in the map illustrate the land cover type. **B.** Heatmap of physicochemical properties of each soil sample. Dendrogram shows the grouping of samples based on these properties. **C.** Principal component analysis of physicochemical properties of soil samples.

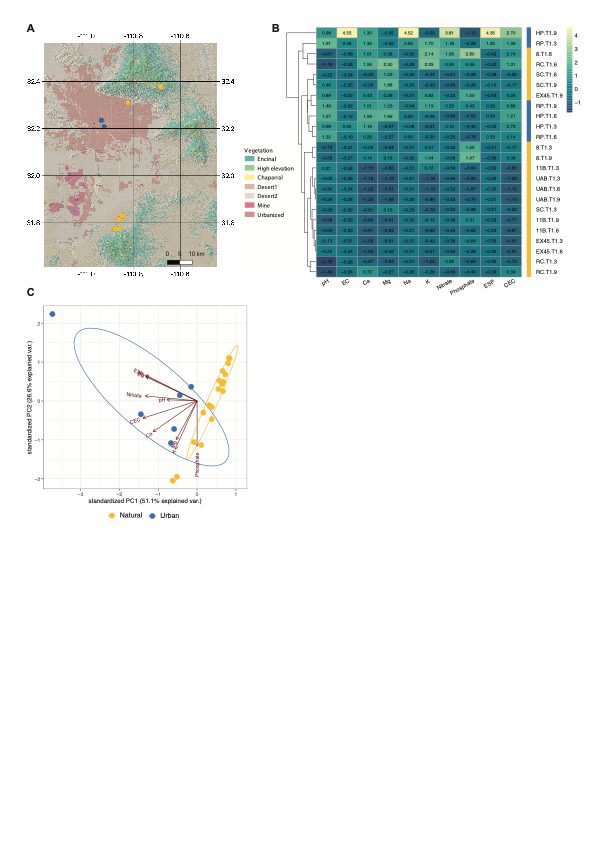

**Supplementary Figure 4. Bacterial functional groups and viral abundances. A., B . and C.** Bacterial functional traits in natural and urban soils (Variance of GC content: X^2^ = 3.69, p-value = 0.055; Codon usage bias: X^2^ = 0.33, p-value = 0.565; Proportion of unannotated genes: X^2^ = 0.40, p-value = 0.525). **F. and E.** Boxplot showing the abundance in RPKM of viruses in natural and urban soils, and colored by vegetation.

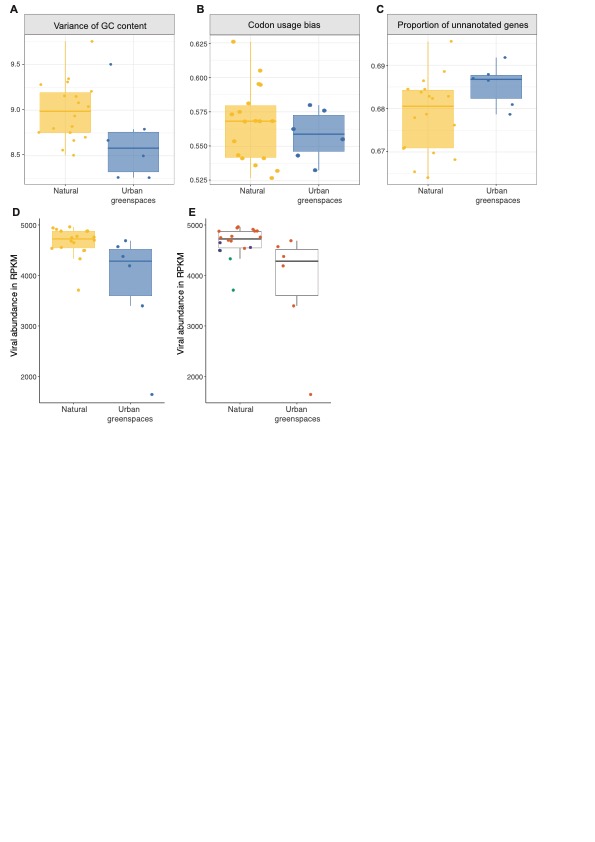

**Supplementary figure 5. Viral and host taxonomy. A. and B.** Taxonomy of predicted viral hosts at the class and family level respectively. **C.** Viral taxonomy at the family level when possible, and at broader groups level when other information was not possible.

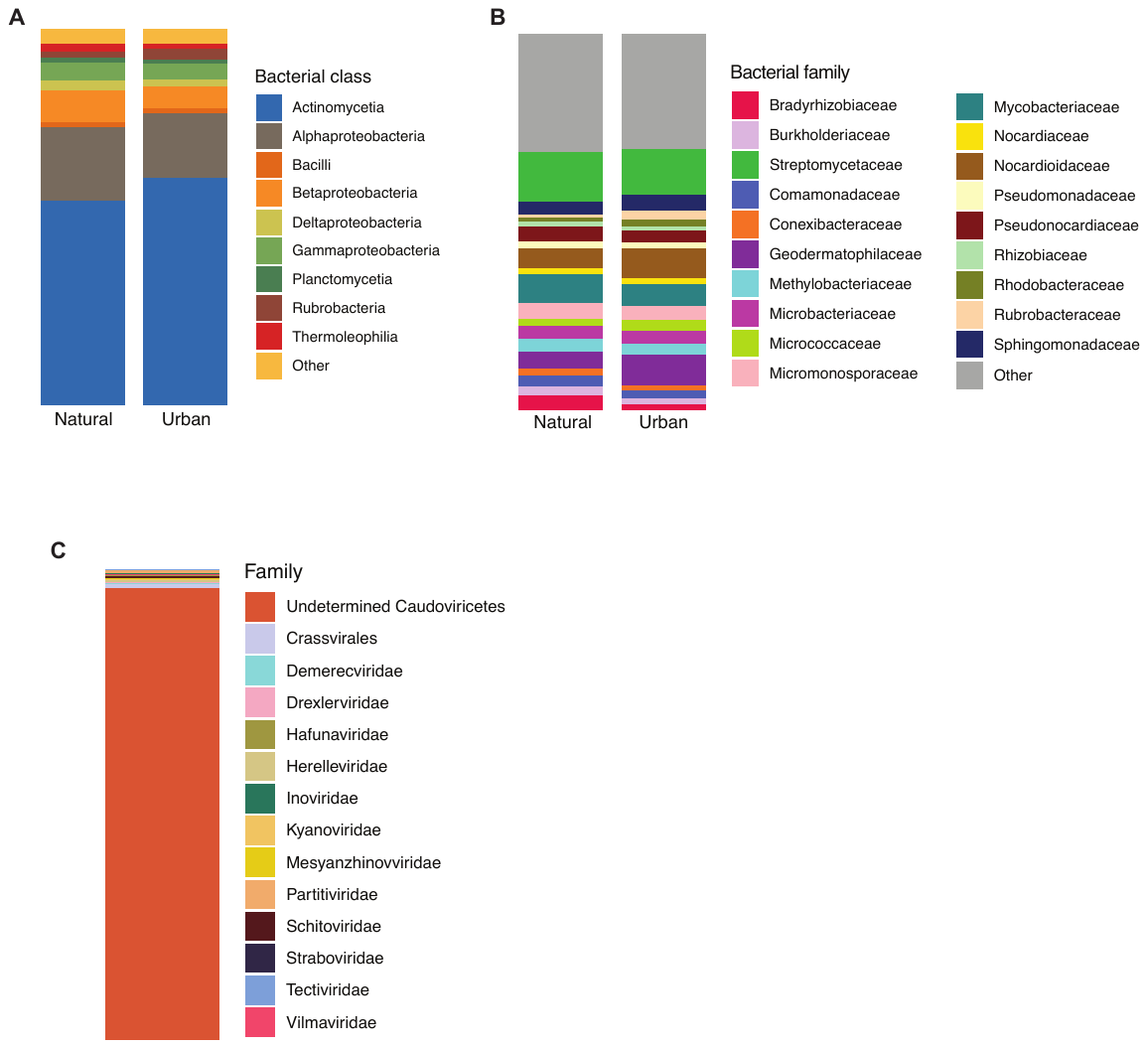

**Supplementary Figure 6. Virus host relationships in urban greenspace and natural soils. A-D,** Density distribution of Pearson correlation coefficient values. **A-B**, Correlation was calculated between the virus-host relative abundance ratio (VHR) and the abundance of the host. **C-D,** Relationship between the abundance of a bacterial host in a sample and the number of virus that target said host (putative viral link) in that same sample.

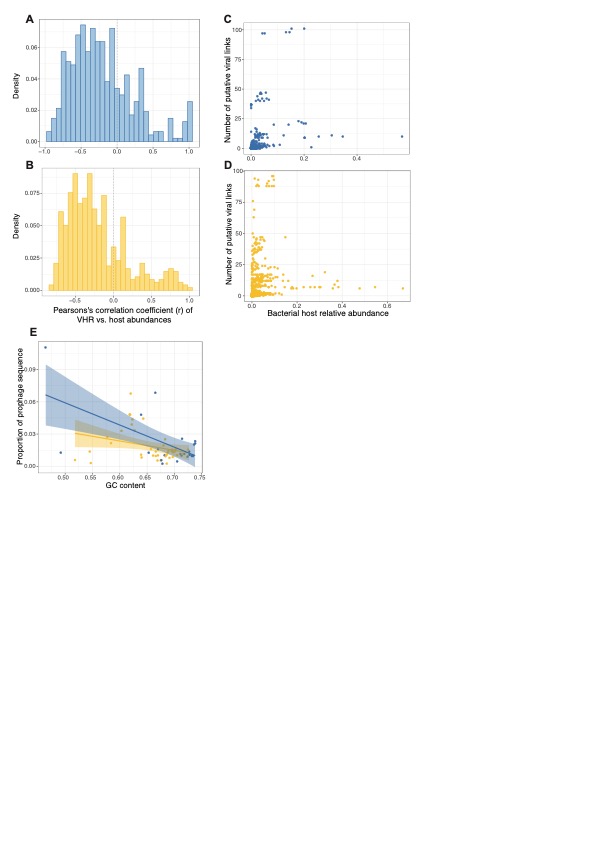

**Supplementary Figure 7.** Relative abundance of auxiliary metabolic genes found in viral genomes.
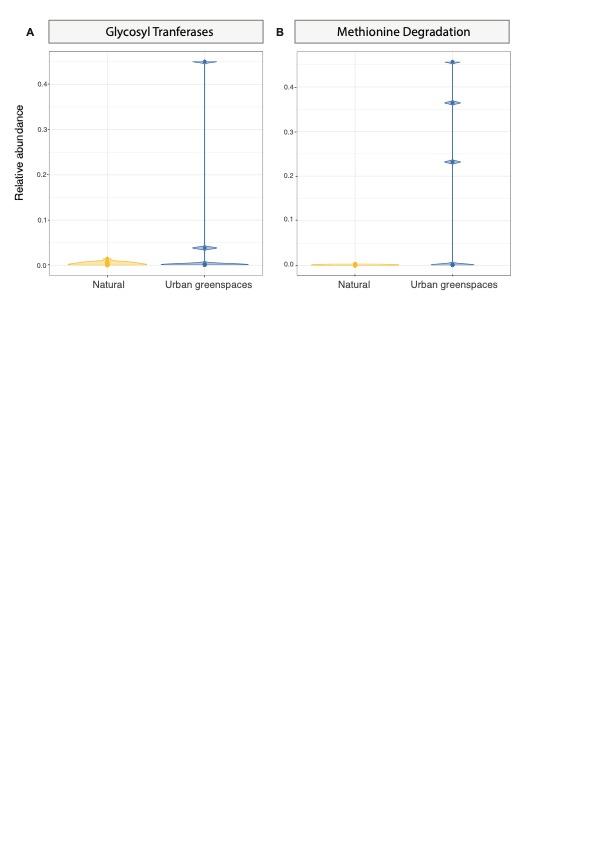

**Supplementary figure 8. Diversity and abundance of carbon and nitrogen cycling genes based on vegetation and denitrification breakdown. A. and B.** Abundance in RPKM and number of different carbon cycling genes based on vegetation type. **C and D.** Abundance in RPKM and number of different nitrogen cycling genes based on vegetation type. **E.** Representation of the steps in denitrification and abundance in RPKM of genes involved in denitrification in natural and urban greenspace soils. **F.** Abundance in RPKM of the genes involved in ammonia oxidation.

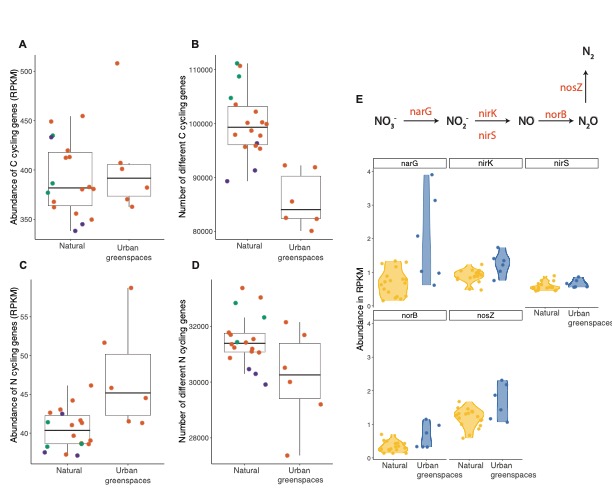

**Supplementary Figure 9. Diversity and abundance of heavy metal and antibiotic resistance genes based on vegetation and denitrification breakdown. A. and B.** Abundance in RPKM and number of different heavy metal resistance genes based on vegetation type. **C and D.** Abundance in RPKM and number of different antibiotic resistance genes based on vegetation type.

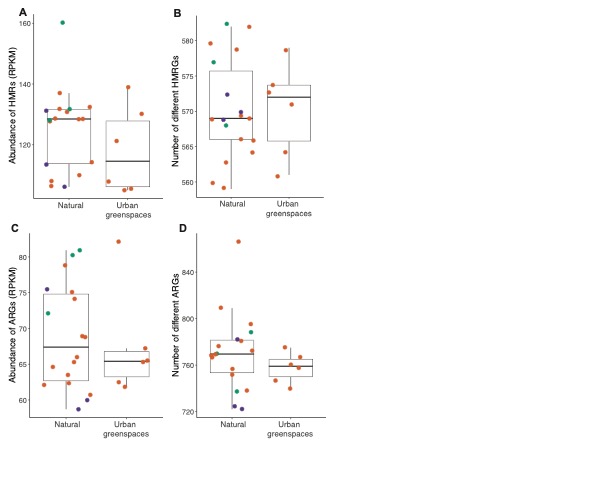

**Supplementary Figure 10. Differential abundance of genes of resistance to different heavy metals and antibiotic classes and pathways. A.** Differential abundance of genes of resistance to different heavy metals between urban greenspace and natural soils. **B.** Differential abundance of genes of resistance to different antibiotic classes between urban greenspace and natural soils. **C.** Abundance in RPKM of antibiotic resistance genes with different mechanisms.

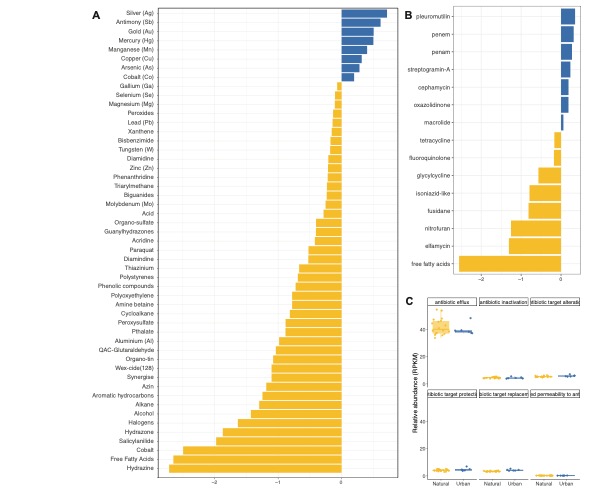

**Supplementary Figure 11. MAGs taxonomy.**

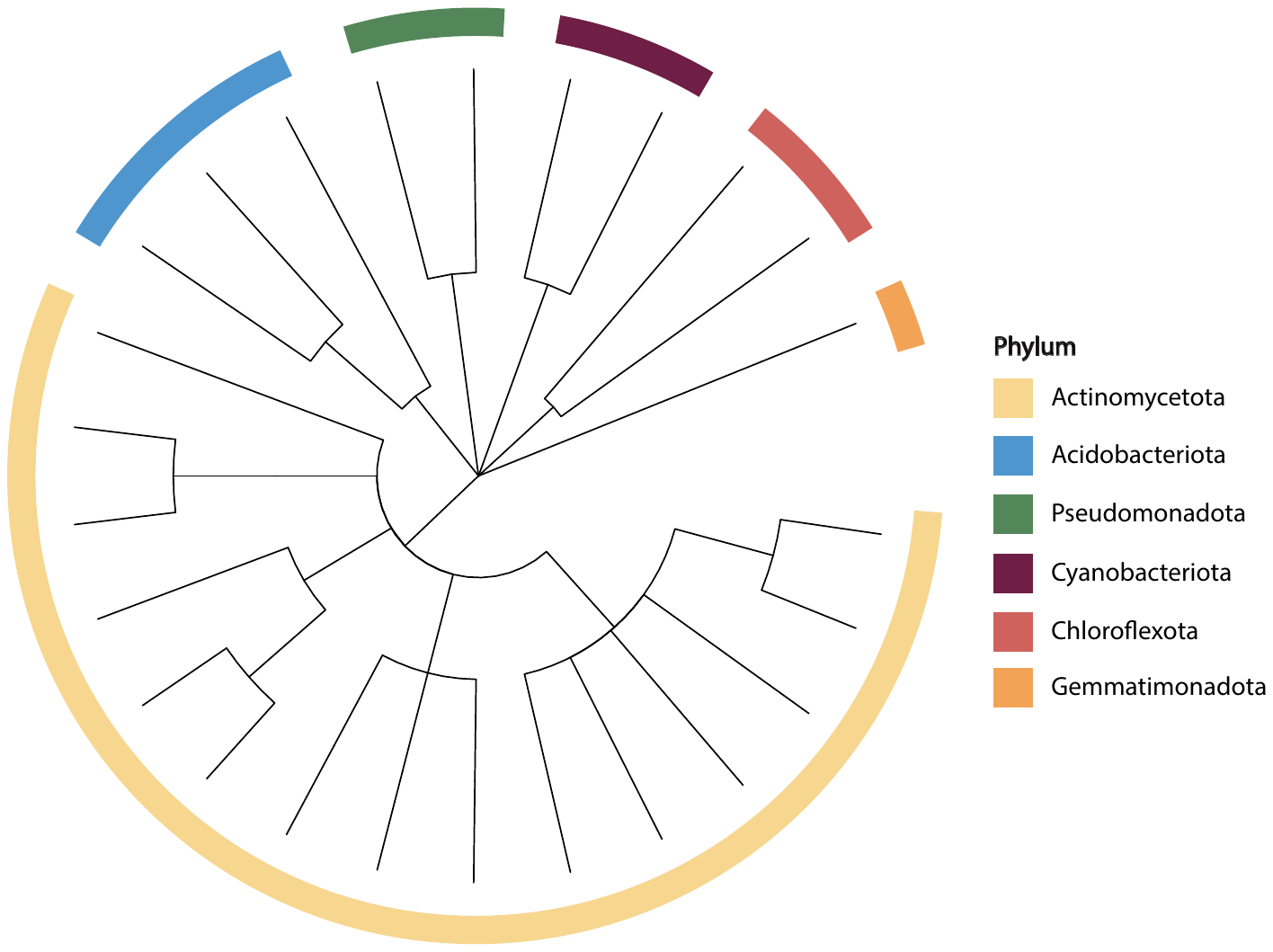

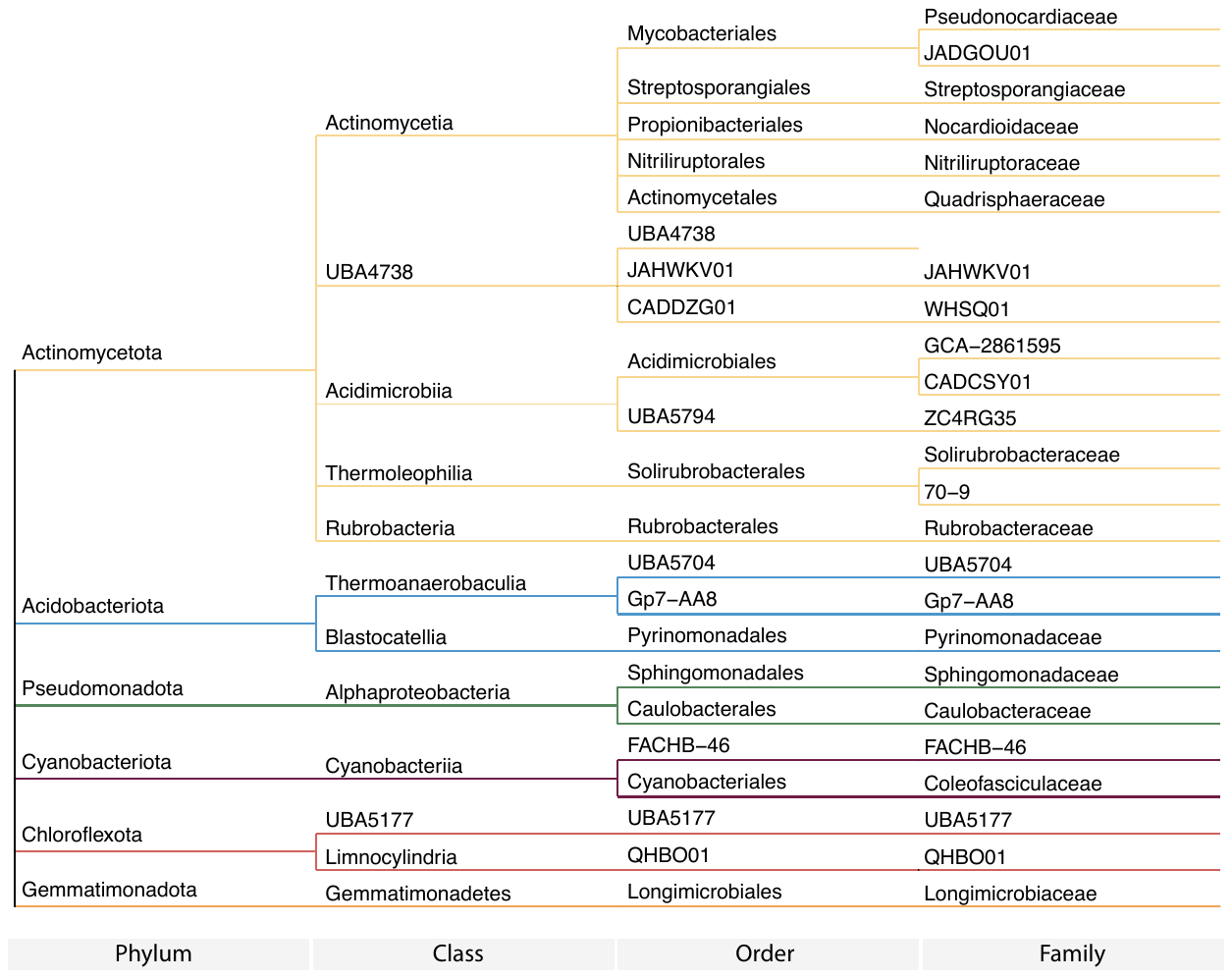

**Supplementary Figure 12. MAGs abundance per sample and genetic content.**

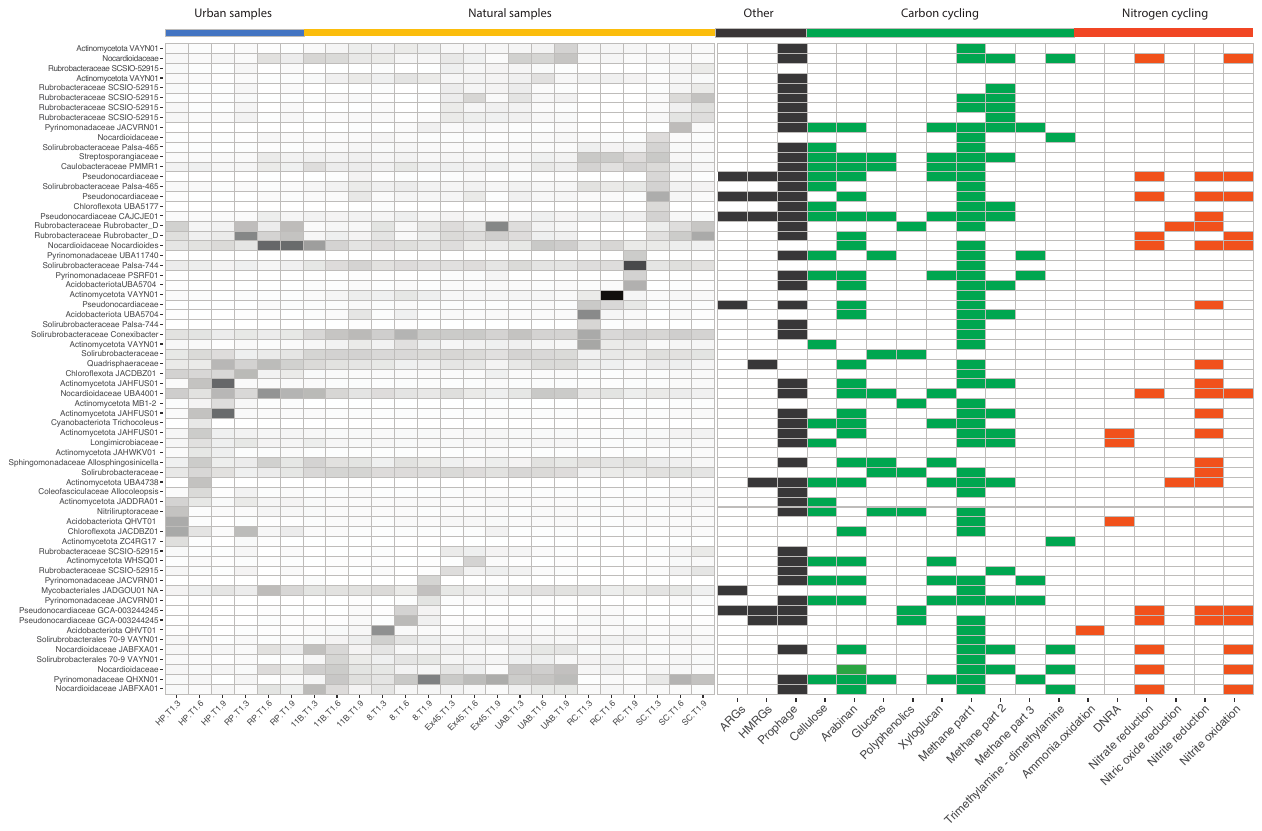

**Table 1. Soil chemical properties**

|  | **pH** | **EC** | **Ca** | **Mg** | **Na** | **K** | **NO_3-_** | **PO_4_^3-^** | **ESP** | **CEC** | **Urban/Natural** |
| --- | --- | --- | --- | --- | --- | --- | --- | --- | --- | --- | --- |
| **HP.T1.3** | 7.9 | 1.4 | 2000 | 100 | 120 | 130 | 12 | 15 | 4.5 | 11.7 | Urban greenspace |
| **HP.T1.6** | 8.6 | 0.34 | 2300 | 190 | 160 | 200 | 1.4 | 14 | 4.9 | 14.3 | Urban greenspace |
| **HP.T1.9** | 7.9 | 7 | 2100 | 110 | 2300 | 140 | 61 | 8.9 | 45.9 | 21.8 | Urban greenspace |
| **RP.T1.3** | 9 | 1.2 | 2100 | 110 | 610 | 440 | 26 | 16 | 17.5 | 15.2 | Urban greenspace |
| **RP.T1.6** | 8.3 | 0.4 | 1400 | 100 | 160 | 160 | 6.8 | 12 | 7.8 | 8.9 | Urban greenspace |
| **RP.T1.9** | 8.5 | 0.52 | 1900 | 170 | 140 | 370 | 14 | 22 | 4.9 | 12.5 | Urban greenspace |
| **RC.T1.3** | 4.7 | 0.15 | 740 | 73 | 14 | 69 | 14 | 13 | 1.3 | 4.5 | Natural |
| **RC.T1.6** | 5.4 | 0.2 | 2300 | 220 | 24 | 520 | 18 | 23 | 0.7 | 14.8 | Natural |
| **RC.T1.9** | 5 | 0.06 | 1700 | 100 | 22 | 170 | 1.4 | 15 | 1 | 9.9 | Natural |
| **SC.T1.3** | 6.3 | 0.1 | 780 | 120 | 20 | 100 | 2.2 | 16 | 1.7 | 5.2 | Natural |
| **SC.T1.6** | 6.5 | 0.06 | 910 | 170 | 16 | 150 | 2.1 | 16 | 1.1 | 6.4 | Natural |
| **SC.T1.9** | 7.3 | 0.05 | 1000 | 200 | 63 | 150 | 1 | 16 | 3.7 | 7.3 | Natural |
| **UAB.T1.3** | 6.7 | 0.03 | 400 | 60 | 12 | 50 | 1.3 | 10 | 1.9 | 2.7 | Natural |
| **UAB.T1.6** | 6.4 | 0.06 | 360 | 56 | 9.9 | 58 | 3.3 | 11 | 1.8 | 2.5 | Natural |
| **UAB.T1.9** | 6.3 | 0.06 | 350 | 62 | 16 | 91 | 3.3 | 12 | 2.7 | 2.6 | Natural |
| **11B.T1.3** | 7 | 0.2 | 420 | 74 | 14 | 230 | 5.6 | 13 | 1.8 | 3.4 | Natural |
| **11B.T1.6** | 6 | 0.08 | 570 | 69 | 12 | 120 | 3.6 | 14 | 1.4 | 3.8 | Natural |
| **11B.T1.9** | 5.6 | 0.09 | 630 | 70 | 15 | 190 | 5.4 | 21 | 1.5 | 4.3 | Natural |
| **8.T1.3** | 5.9 | 0.11 | 1200 | 87 | 14 | 210 | 5.9 | 31 | 0.8 | 7.3 | Natural |
| **8.T1.6** | 5.8 | 0.44 | 1900 | 130 | 18 | 500 | 31 | 42 | 0.7 | 11.9 | Natural |
| **8.T1.9** | 6.1 | 0.16 | 1300 | 120 | 19 | 350 | 9.5 | 35 | 1 | 8.5 | Natural |
| **EX45.T1.3** | 6.6 | 0.11 | 490 | 84 | 11 | 150 | 5 | 18 | 1.3 | 3.6 | Natural |
| **EX45.T1.6** | 6.5 | 0.07 | 480 | 93 | 11 | 130 | 3.3 | 16 | 1.3 | 3.6 | Natural |
| **EX45.T1.9** | 7.8 | 0.24 | 1500 | 130 | 12 | 320 | 7.2 | 31 | 0.6 | 9.5 | Natural |
| **Average** |  |  |  |  |  |  |  |  |  |  |  |
|  | **pH** | **EC** | **Ca** | **Mg** | **Na** | **K** | **NO_3-_** | **PO_4_^3-^** | **ESP** | **CEC** |  |
| **Urban** | 8.37 | 1.81 | 1966.67 | 130 | 581.67 | 240 | 20.2 | 14.65 | 14.25 | 14.07 |  |
| **Natural** | 6.22 | 0.13 | 946.11 | 106.56 | 17.94 | 197.67 | 6.84 | 19.61 | 1.46 | 6.21 |  |

**Table 2. List of all inferred auxiliar metabolic genes (AMGs).**

| **gene** | **gene_description** | **module** | **sheet** | **reference** |
| --- | --- | --- | --- | --- |
| HP-T1-6_contig.fa_viral_101__full-cat_2_4 | dCTP deaminase [EC:3.5.4.13] [RN:R02325] | Pyrimidine deoxyribonuleotide biosynthesis | MISC |  |
| HP-T1-6_contig.fa_viral_242__full-cat_2_3 | ferredoxin | Cytochrome b6f complex | Energy |  |
| HP-T1-6_contig.fa_viral_288__full-cat_1_6 | nuoD; NADH-quinone oxidoreductase subunit D [EC:7.1.1.2] | NADH:quinone oxidoreductase, prokaryotes | Energy |  |
| HP-T1-6_contig.fa_viral_47__full-cat_2_39 | DNA (cytosine-5-)-methyltransferase [EC:2.1.1.37] [RN:R04858] | Methionine degradation | Organic Nitrogen |  |
| HP-T1-9_contig.fa_viral_101__full-cat_2_16 | DNA (cytosine-5-)-methyltransferase [EC:2.1.1.37] [RN:R04858] | Methionine degradation | Organic Nitrogen |  |
| HP-T1-9_contig.fa_viral_174__full-cat_2_14 | DNA (cytosine-5-)-methyltransferase [EC:2.1.1.37] [RN:R04858] | Methionine degradation | Organic Nitrogen |  |
| RP-T1-3_contig.fa_viral_659__full-cat_2_10 | DNA (cytosine-5-)-methyltransferase [EC:2.1.1.37] [RN:R04858] | Methionine degradation | Organic Nitrogen |  |
| RP-T1-3_contig.fa_viral_779__full-cat_2_3 | DNA (cytosine-5-)-methyltransferase [EC:2.1.1.37] [RN:R04858] | Methionine degradation | Organic Nitrogen |  |
| RP-T1-3_contig.fa_viral_838__full-cat_1_8 | GT2 cellulose synthase* | GlycosylTransferases | carbon utilization |  |
| RP-T1-6_contig.fa_viral_476__full-cat_1_2 | 1,4-dihydroxy-2-naphthoyl-CoA hydrolase | Menaquinone biosynthesis | MISC |  |
| RP-T1-9_contig.fa_viral_121__full-cat_1_5 | 1,4-dihydroxy-2-naphthoyl-CoA hydrolase | Menaquinone biosynthesis | MISC |  |
| HP-T1-6_contig.fa_viral_242__full-cat_2_3 | petE; ferredoxin | M00161 |  | Sullivan et al. 2005; Doron et al. 2016 |
| HP-T1-6_contig.fa_viral_242__full-cat_2_3 | petF; ferredoxin | M00161 |  | Sullivan et al. 2005 |
| HP-T1-9_contig.fa_viral_112__full-cat_2_9 | Concanavalin A-like lectin; extracellular arabinase |  |  | Emerson et al. 2018 |
| HP-T1-9_contig.fa_viral_48__full-cat_1_10 | Sulfotransferase family |  |  | Roux et al. 2016 |
| RC-T1-3_contig.fa_viral_40__full-cat_2_8 | 6-pyruvoyl tetrahydropterin synthase |  |  | Roux et al. 2016 |
| RP-T1-3_contig.fa_viral_838__full-cat_1_9 | O-methyltransferase |  |  | Roux et al. 2016 |
| UAB-T1-6_contig.fa_viral_115__full-cat_1_28 | Concanavalin A-like lectin; extracellular arabinase |  |  | Emerson et al. 2018 |

* (EC 2.4.1.12); chitin synthase (EC 2.4.1.16); dolichyl-phosphate beta-D-mannosyltransferase (EC 2.4.1.83); dolichyl-phosphate beta-glucosyltransferase (EC 2.4.1.117); N-acetylglucosaminyltransferase (EC 2.4.1.-); N-acetylgalactosaminyltransferase (EC 2.4.1.-); hyaluronan synthase (EC 2.4.1.212); chitin oligosaccharide synthase (EC 2.4.1.-); beta-1,3-glucan synthase (EC 2.4.1.34); beta-1,4-mannan synthase (EC 2.4.1.-); beta-mannosylphosphodecaprenol-mannooligosaccharide alpha-1,6-mannosyltransferase (EC 2.4.1.199); UDP-Galf: rhamnopyranosyl-N-acetylglucosaminyl-PP-decaprenol beta-1,4/1,5-galactofuranosyltransferase (EC 2.4.1.287); UDP-Galf: galactofuranosyl-galactofuranosyl-rhamnosyl-N-acetylglucosaminyl-PP-decaprenol beta-1,5/1,6-galactofuranosyltransferase (EC 2.4.1.288); dTDP-L-Rha: N-acetylglucosaminyl-PP-decaprenol alpha-1,3-L-rhamnosyltransferase (EC 2.4.1.289)
